# Metagenomic identification of diverse animal hepaciviruses and pegiviruses

**DOI:** 10.1101/2020.05.16.100149

**Authors:** Ashleigh F. Porter, John H.-O. Pettersson, Wei-Shan Chang, Erin Harvey, Karrie Rose, Mang Shi, John-Sebastian Eden, Jan Buchmann, Craig Moritz, Edward C. Holmes

## Abstract

The RNA virus family *Flaviviridae* harbours several important pathogens of humans and other animals, including Zika virus, dengue virus and hepatitis C virus. The *Flaviviridae* are currently divided into four genera - *Hepacivirus*, *Pegivirus*, *Pestivirus* and *Flavivirus* – each of which have a diverse host range. Members of the genus *Hepacivirus* are associated with a diverse array of animal species, including humans and non-human primates, other mammalian species, as well as birds and fish, while the closely related pegiviruses have been identified in a variety of mammalian taxa including humans. Using a combination of meta-transcriptomic and whole genome sequencing we identified four novel hepaciviruses and one novel variant of a known virus, in five species of native Australian wildlife, expanding our knowledge of the diversity in this important group of RNA viruses. The infected hosts comprised native Australian marsupials and birds, as well as a native gecko (*Gehyra lauta*). The addition of these novel viruses led to the identification of a distinct marsupial clade within the hepacivirus phylogeny that also included an engorged *Ixodes holocyclus* tick collected while feeding on Australian long-nosed bandicoots (*Perameles nasuta*). Gecko and avian associated hepacivirus lineages were also identified. In addition, by mining the short-read archive (SRA) database we identified another five novel members of *Flaviviridae*, comprising three new hepaciviruses from avian and primate hosts, as well as two primate-associated pegiviruses. The large-scale phylogenetic analysis of these novel hepacivirus and pegivirus genomes provides additional support for virus-host co-divergence over evolutionary time-scales.

## Introduction

As the vast majority of emerging infectious disease in humans are caused by viral zoonoses [1], the identification and characterization of animal viruses is critical for identifying potential disease reservoirs and providing models for the study of human viruses [2,3]. Two related groups of viruses that have recently received considerable attention are the genera *Hepacivirus* and *Pegivirus* from the family *Flaviviridae* of single-strand positive-sense RNA viruses. Hepaciviruses infect a broad range of vertebrate hosts, including humans, non-human primates [4,5], and a variety of other mammals including rodents [6–10], horses [11], bats [12], and cows [13,14]. Hepaciviruses have also been detected in birds [15, 16], fish and a variety of other vertebrates [17, 18]. Additionally, two hepacivirus-like sequences have been identified in arthropods (a mosquito and tick), although their true host is uncertain [19, 20]. Despite such a wide diversity of animal hosts, hepaciviruses remain synonymous with liver infection, with the most notable example being human hepatitis C virus (HCV). While some non-primate hepaciviruses have been well characterized, particularly equine hepacivirus (also known as Hepacivirus AK, or non-primate hepacivirus, NPHV) and canine hepacivirus (CHV), it seems likely that HCV, along with NPHV and CHV, arose from a currently unknown zoonotic source, with rodents and bats suggested as possible reservoirs [21–24].

The genus *Pegivirus* contains 11 defined virus species known to infect humans and variety of other mammals. In addition, a novel avian pegivirus was recently identified in a Common myna bird (*Acridotheres tristis*) [25]. Human pegivirus (HPgV), previously known as GB virus C, is known to infect humans but with an unproven link with clinical illness [26], despite being identified in the brain tissue of patients with encephalitis [27]. Non-human primate pegiviruses have been identified in both New World monkeys [28, 29] and Old World apes, including chimpanzees [30–32], the latter of which (SPgV_cpz_) are closely related to HPgV [33]. Other pegiviruses have been found in horses [34], bats [12, 35] and rodents [7,8].

Despite their broad host range, all hepaciviruses and pegiviruses described to date possess a single-strand positive-sense RNA genome and a large open reading frame encoding a single polyprotein flanked by untranslated regions. Although this structure is common among the *Flaviviridae* [36, 37], a more diverse set of genome structures, including segmented forms, have been identified in invertebrate flaviviruses [18]. The multifunctional polyprotein is cleaved by proteases to create ten proteins: three structural (core, E1, E2) and seven non-structural (NS1, NS2, NS3, NS4a, NS4b, NS5a, NS5b) [24]. It is important to note that while many members of the genus *Flavivirus* are transmitted via arthropod vectors, and it has been hypothesised that biting arthropod vectors may transmit some hepaciviruses [37], there is currently no evidence of vector-borne transmission in either the *Hepacivirus* or *Pegivirus* genera. Recently, however, a novel hepacivirus, Brushtail possum hepacivirus, was discovered in brush-tailed possums (*Trichosurus vulpecula*), a native Australian marsupial [38]. Similarly, Collins beach hepacivirus was identified in a tick (*Ixodes holocyclus*) feeding on another Australian marsupial (long-nosed bandicoot, *Perameles nasuta*) [19]. These data suggest that Australian native wildlife harbour a diversity of hepaciviruses worthy of further exploration.

As hepaciviruses have traditionally been difficult to culture, the recent expansion of animal hepaciviruses is largely due to the advent of unbiased high-throughput sequencing. This expansion has greatly impacted our understanding of the diversity and evolution of this important genus. Herein, we used a bulk RNA-sequencing (meta-transcriptomic) approach to identify additional novel hepaciviruses in Australian wildlife and determine their abundance. This approach has previously proven successful in an Australian context, identifying a variety of viruses in Australian wildlife, including native Australian and invasive species [19, 38–43]. To supplement this analysis, we mined the sequence read archive (SRA) database for hepacivirus-like sequences. Previously generated RNA sequencing data, such as those found in the SRA, are a relatively untapped resource for novel virus discovery [44]. Through SRA-mining we identified three novel hepaciviruses and two novel pegiviruses, again highlighting the power of genomics to identify novel viral sequences.

## Materials and Methods

### Animal ethics

The magpie lark and pelican were handled under a series of NSW Office of Environment and Heritage Licences to Rehabilitate Injured, Sick or Orphaned Protected Wildlife (#MWL000100542). Samples were collected under the Opportunistic Sample Collection Program of the Taronga Animal Ethics Committee, and scientific licence SL100104 issued by the NSW Office of Environment and Heritage. Ticks were removed from the long-nosed bandicoot under the approval of the NSW Office of Environment and Heritage Animal Ethics Committee (#000214/05) and scientific license SL100104.

### Sample collection

The samples analysed here were collected from a variety of sources (Table 1). Gecko hepacivirus (RNASeq library VERT7) was obtained from a liver sample (#CCM0247) of a seemingly healthy gecko (*Gehyra lauta*) collected at West Leichardt Station, Queensland in 2013 [45]. Pelican hepacivirus (RNASeq library VERT5) was obtained from the liver (Australian Registry of Wildlife Health #9381.1) of an Australian pelican (*Pelecanus conspicillatus*) collected at Blackwall Bay, New South Wales (NSW) in 2013. The pelican presented with an ongoing syndrome of profound weakness, diarrhoea, dyspnoea, and death, consistently associated with myocardial degeneration to necrosis. Magpie lark hepacivirus (Australian Registry of Wildlife Health #9585.8) was obtained from brain (RNASeq library VERT14) and heart samples (RNASeq library VERT 15) of an injured juvenile magpie lark (*Grallina cyanoleuca*) collected at Warigee, NSW in 2013 [46]. Collins Beach virus 1 was identified from three engorged female ticks (*Ixodes holocylus)* collected while feeding on a healthy long-nose bandicoot (*Perameles nasuta*) at North Head, NSW in 2016. These individual tick samples were part of a larger pool of ticks (RNASeq library TICK08, SRA projects SRS3932533 and SRS3932534) used to identify the partial genome sequence of Collins Beach virus [19]. The Koala hepacivirus was identified during our initial SRA mining of marsupial transcriptomes (Supplementary Table 1). In this case, the mRNA library (SRX501262) was prepared from the liver RNA of a deceased Australian koala (*Phascolarctos cinereus)* “Pacific Chocolate” that was known to be infected with chlamydia [46]. As viral coverage was incomplete due to the poly-A selection method, the original tissue sample (#M.45022.004) was kindly provided by the Australian Museum and subjected to total RNA sequencing (i.e. rRNA-depletion only) along with our other cases.

**Table 1.**
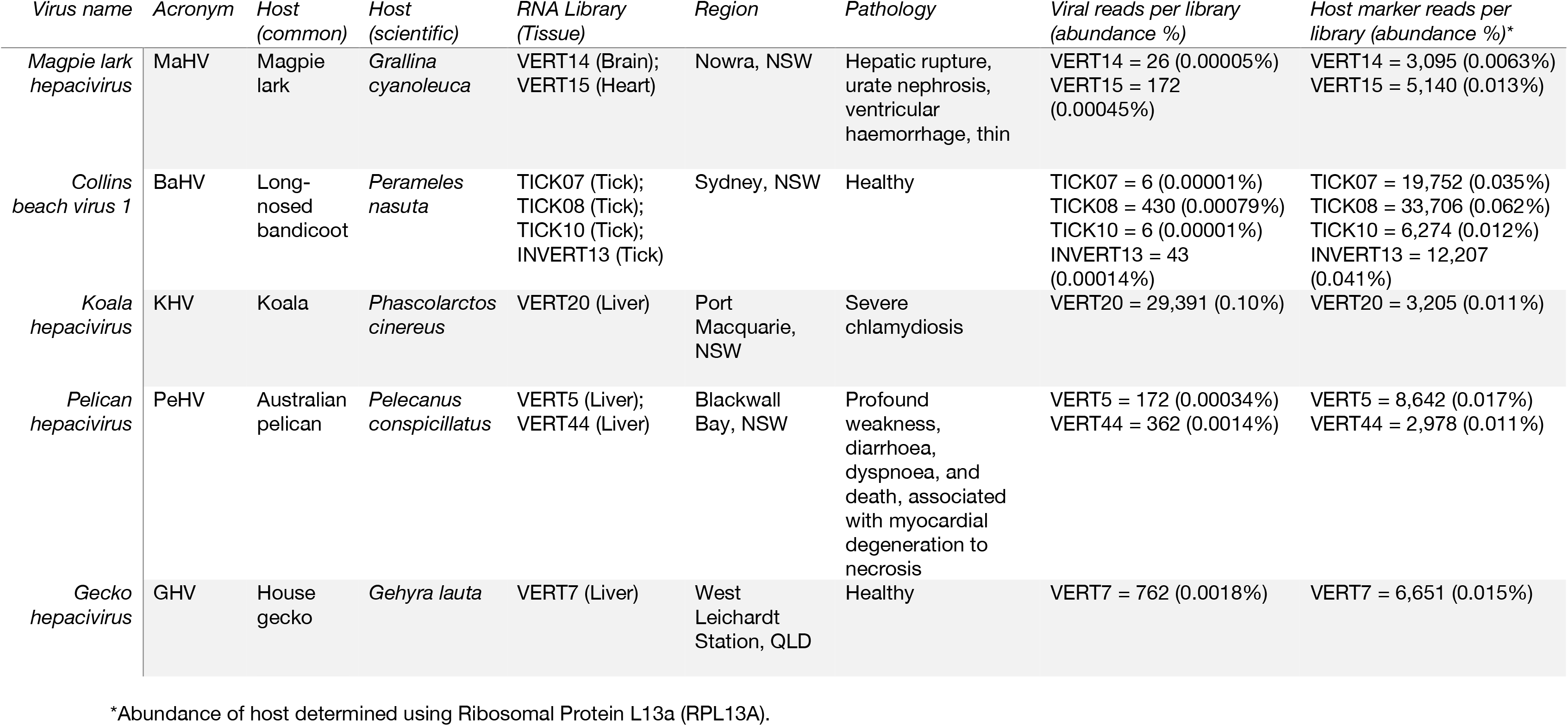
Sample information for the novel hepaciviruses identified here, including the host species, library, region isolated, host pathology, and assembly data.

### RNA extraction, library preparation and sequencing

Viral RNA was extracted from individual tissues of animal samples using the RNeasy Plus Mini Kit (Qiagen, Germany). RNA concentration and integrity were determined using a NanoDrop spectrophotometer (ThermoFisher) and TapeStation (Agilent). RNA samples were pooled in equal proportions based on animal tissue types and syndrome (maximum eight individuals per library). Illumina TruSeq stranded RNA libraries were prepared on the pooled samples following rRNA depletion using a RiboZero Gold kit (Epidemiology) at the Australian Genome Research Facility (AGRF), Melbourne. The rRNA depleted libraries were then sequenced on an Illumina HiSeq 2500 system (Paired end 100 bp or 125 bp reads) to depths of between 13-28 million paired reads.

### Viral discovery pipeline

We employed an established meta-transcriptomic pipeline developed for RNA virus discovery [18, 19, 47]. RNA sequences were trimmed of low-quality bases and adapter sequences with Trimmomatic v0.36 [48] before *de novo* assembly using Trinity 2.5.1 [49]. The assembled contigs were then compared against the NCBI nucleotide (nt) and non-redundant (nr) protein databases with Blastn v2.7.1+ [50] and Diamond v0.9.18 [51], respectively, with an e-value threshold of 1E-5. Contig abundance was estimated by calculating the number of reads in each library that mapped to the hepacivirus genome divided by the number of total reads (Supplementary Table 2). Similarly, host abundance was compared by mapping reads to a reference gene - ribosomal protein L13a (RPL13A). Low abundance viruses (i.e. those that could not be assembled) were identified by aligning the trimmed reads against either NCBI viral RefSeq or a curated hepacivirus/pegivirus protein database using Diamond v.0.9.18, with an e-value threshold of 1E-4. Curated hepacivirus/pegivirus databases were regularly updated with novel viruses to increase read alignment for divergent viral species.

### PCR confirmation and whole-genome sequencing

To confirm the presence of novel viruses in individual samples, RT-PCR was performed using primers designed to amplify a short region of the viral genome (<300bp). Briefly, Superscript IV VILO master mix was used to generate cDNA from individual RNA samples before RT-PCR screening with Platinum SuperFi. Following the confirmation of viruses to individual samples, long (2-4kb) overlapping PCRs were designed to amplify the available viral genome (i.e. complete or partial) for the hepaciviruses identified and sequenced using the Nextera XT library prep kit and Illumina MiSeq sequencing (150nt paired reads). Viral genomes were then assembled using MegaHit v1.1.3 [52].

### SRA mining

To identify hepacivirus- and pegivirus-like viruses present in the SRA, we screened primates (excluding *Homo sapiens)*, birds (excluding *Gallus gallus)* and marsupials. We focused on these taxonomic groups because (i) although HCV is clearly a virus of humans, it is unclear whether it is present in other non-human primates (which may be indicative of virus-host co-divergence), (ii) a number of novel hepacivirus- and pegivirus-like viruses have recently been identified in avian hosts suggesting that these might be a rich source of novel viruses, and (iii) to confirm the presence of a marsupial specific-lineage as suggested by our previous studies. Accordingly, three different sets of single- and paired-end RNA sequencing SRAs were analysed: one included 2,312 avian SRAs (Supplementary Table 3), a second that included 3,902 primate data sets (Supplementary Table 4) and finally a marsupial data set of 330 SRAs (Supplementary Table 1). SRAs were fetched with the NCBI SRA toolkit using fastq-dump and analysed using our virus discovery pipeline, with the exception that the initial viral screening was performed by aligning the downloaded reads to our curated hepacivirus/pegivirus protein database with Diamond v.0.9.18. SRAs that contained hepacivirus-like and/or pegivirus-like reads were then assembled and annotated as above.

Three virus fragments were identified in three runs (SRR1325073, SRR1325072, SRR1325074) from SRA project SRX565268, isolated from Eurasian blue tit (*Cyanistes caeruleus)* from Germany. These three fragments were concatenated into a single genome, Blue tit hepacivirus (BtHV). The three fragments, and their respective coding regions, are shown in Supplementary Figure 1.

### Genome annotation

The hepaciviruses identified from our samples and via SRA mining were subjected to an online sequence similarity search using NCBI blastx as well as a conserved domain (CDD) search. ORFs of verified contigs were included in a phylogenetic analysis as described below. Genome annotation of the 10 novel virus sequences was performed using Geneious Prime (version 2019.2.1) [53]. Specifically, using the Live Annotate and Predict tool, each novel hepacivirus genome was screened for hepacivirus-specific gene annotations utilising previously identified annotations in hepacivirus genomes downloaded from NCBI (n = 145).

To maximise the identification of hypothetical genes, even in divergent genomes, the gene similarity cut-off was set to 25%, with predicted genes then manually annotated.

### Phylogenetic analysis

To investigate the evolutionary relationships among the hepaciviruses and pegiviruses we collated the amino acid sequences for the complete polyprotein of each genome from the ten novel viruses identified here (see Results), and from hepaciviruses and pegiviruses taken from NCBI Refseq database including Brushtail possum hepacivirus and Collins Beach hepacivirus (n =110, accession numbers available in Supplementary Table 5). These sequences were aligned using the E-INS-i algorithm in MAFFT v7 [54], with ambiguous regions removed using GBlocks [55]. The final sequence alignment comprised 110 sequences of 1345 amino acid residues in length. ProtTest v3.4 [56] was used to determine the most appropriate model of amino acid substitution for these data. From this, we estimated a phylogenetic tree using the maximum likelihood method in IQ-TREE, version 1.6.12 [57], employing the LG model of amino acid substitution with invariable sites and gamma model (4 categories), along with a bootstrap resampling analysis using 1000 replicates. Two additional phylogenetic trees were estimated using the NS2/3 (549 amino acid residues) and NS5 (407 amino acid residues) regions using the same procedure as described above (although with 100 bootstrap replicates).

## Results

### Novel hepaciviruses discovered in Australian wildlife

We identified four novel hepaciviruses and one new hepacivirus variant from five meta-transcriptomic libraries obtained from a variety of Australian wildlife samples (Table 1). The four novel hepaci-like genomes comprised: (i) Koala hepacivirus (denoted KHV); (ii) Pelican hepacivirus (PeHV), (ii) Magpie lark hepacivirus (MaHV); and (iv) Gecko hepacivirus (GHV). In addition, we identified a variant of the previously identified Collins beach virus isolated from ticks feeding on long-nosed bandicoots [19]. In accordance with the naming convention for hepacivirus variants [58] we term this Collins beach virus 1 (CBV1). The relative abundance of hepacivirus-like sequences in each library was low (Figure 1 and Table 1), with the abundance of the host RPL13A gene shown for comparison (Figure 1). The Koala hepacivirus was the most abundant, comprising 0.1% of the total reads and a complete genome could be assembled at a mean coverage depth of 386X. Reads associated with CBV1 were at an abundance <1% in all tick libraries. The abundance levels for the remaining novel hepaciviruses ranged from 0.0018-0.00001% of the reads in their respective libraries, and most of the genomes were in-complete. Therefore, for these remaining viruses, we used a long, overlapping amplicon-based sequencing approach to fill gaps in the RNASeq data and from this produce complete or near-complete genomes for further analysis.

**Figure 1.**
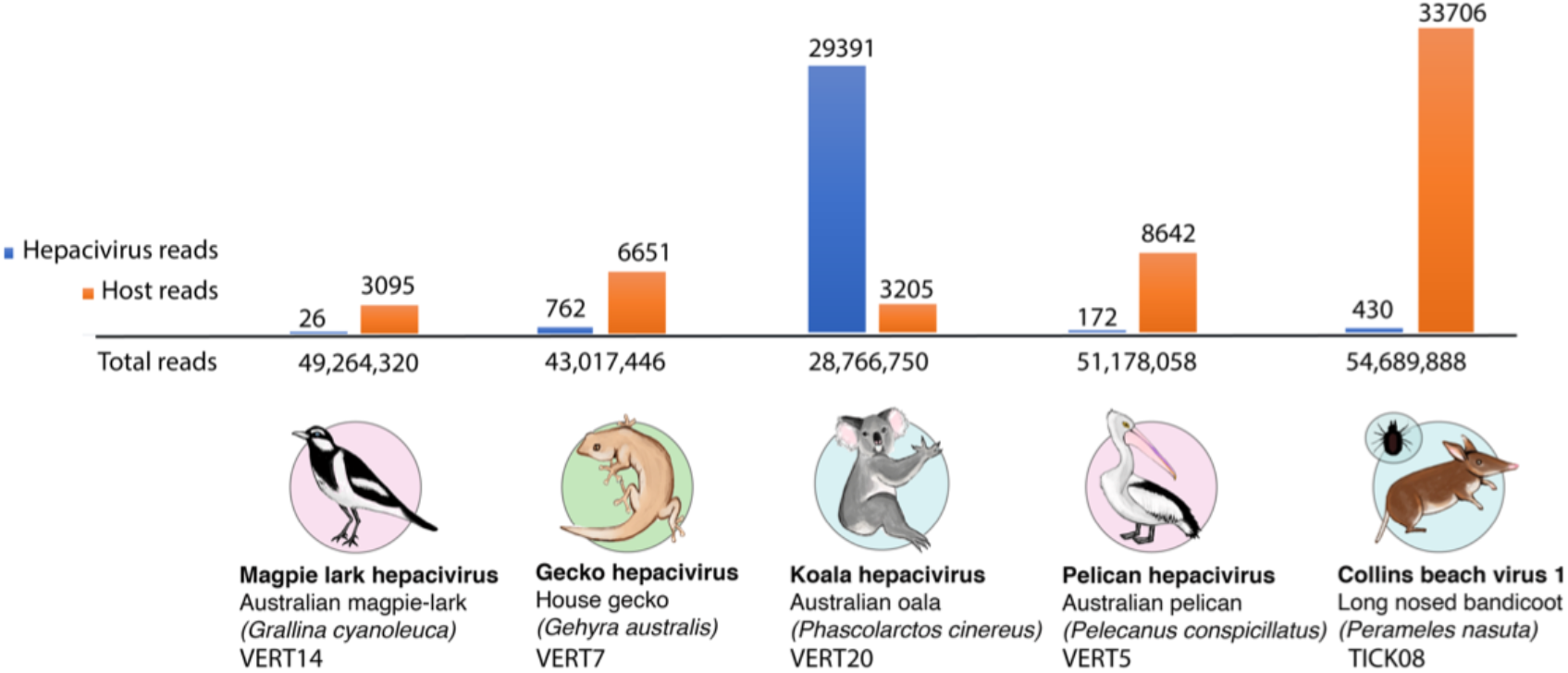
Abundance of hepacivirus-like contigs in each RNA-Seq library. The relative abundance of hepacivirus-like contigs (shown in blue) is presented as the number of hepacivirus-like reads compared to the number of host reads (based on the RPL13A gene, shown in orange) and the total number of reads each library (shown on the horizontal axis).

### Novel hepaciviruses and pegiviruses discovered from SRA data mining

Three hepacivirus sequences and two pegivirus sequences were identified by mining the SRA (Table 2). The hepacivirus genomes identified were isolated from (i) a Diademed sifaka (*Propithecus diadema*) sampled in Toamasina province, Madagascar, and termed Sifaka hepacivirus (SfHV-mad), (ii) a Senegal bushbaby (*Galago senegalensis*) virus termed Bushbaby hepacivirus (BbHV), and (iii) three hepacivirus-like fragments recovered from a Eurasian blue tit (*Cyanistes caeruleus*) in Montpellier, France, that were compiled into a viral single genome termed Blue tit hepacivirus (BtHV). The two pegivirus sequences were isolated from a (i) Common marmoset (*Callithrix jacchus*) termed Marmoset pegivirus (MHV), and (ii) two pegivirus-like fragments recovered from a South African Vervet monkey (*Chlorocebus pygerythrus*) that were assembled into a single viral genome termed Simian pegivirus (SPV-saf). The only marsupial hepacivirus-like sequence identified in the SRA screen was the partial fragment of the Koala hepacivirus identified in library SRX501262 that we re-sequenced to obtain a complete viral genome (VERT20). All individual sequence fragments are described in the Supplementary materials.

**Table 2.** The five novel hepaciviruses and pegiviruses identified from SRA-mining, including host information, project runs (SRR) and SRA project.

| *Virus name* | *Acronym* | *Host*<br>*(common)* | *Host (scientific)* | *Runs (SRR)* | *SRA project* |
| --- | --- | --- | --- | --- | --- |
| *Blue tit*<br>*hepacivirus* | BtHV | Eurasian<br>blue tit | *Cyanistes caeruleus* | SRR1325073,<br>SRR1325072,<br>SRR1325074 | SRX565268 |
| *Bushbaby*<br>*hepacivirus* | BbHV | Senegal<br>bushbaby | *Galago senegalensis* | SRR361358 | SRX104357 |
| *Sifaka*<br>*hepacivirus* | SfHV-mad | Diademed<br>sifaka | *Propithecus diadema* | SRR3131110 | SRX1550742 |
| *Marmoset*<br>*pegivirus* | MPV | Common<br>marmoset | *Callithrix jacchus* | SRR1758976 | SRX843207 |
| *Simian*<br>*pegivirus* | SPV-saf | Vervet<br>monkey | *Chlorocebus*<br>*pygerythrus* | SRR1046735,<br>SRR1046733 | SRX389659 |

### Genome annotation of novel viruses

Each of the novel hepaciviruses identified here, as well as the variant of Collins Beach hepacivirus, underwent genome annotation and were compared (Figure 2) to the fully annotated Bald eagle hepacivirus as a reference [16]. All of the novel viruses encoded near-complete polyproteins (described in Figure 2, highlighted in boxes). Two relatively well conserved *Flaviviridae* proteins, the NS5B protein that contains RNA-dependent RNA polymerase (RdRp) function and the NS3 protease-helicase protein (Figure 2, highlighted in pink and blue, respectively), were present in all the novel flavivirus polyproteins. The remaining flavivirus proteins, comprising the core protein C, protein p7, envelope protein E1, envelope protein E2, NS2, NS4A, NS4b, NS5A were identified in most novel virus genomes, although were unable to successfully identify and annotate NS5A (highlighted in purple, Figure 2) in any of the novel bird or reptile hepaciviruses at our threshold levels (25% sequence similarity). Two of the SRA derived viral sequences - Blue tit hepacivirus and Simian pegivirus - comprised partial genomes only and hence do not include all annotations. The more divergent Gecko hepacivirus was similarly not fully annotated. The genome regions missing from the annotation were the envelope proteins (E1 and E2) and the non-structural proteins NS2 and NS5A (Fig 2). Despite these seemingly incomplete annotations, all of the novel viruses have near-complete polyprotein encoding regions, suggesting that successful annotation has simply been prevented by high levels of sequence divergence in some genes.

**Figure 2.**
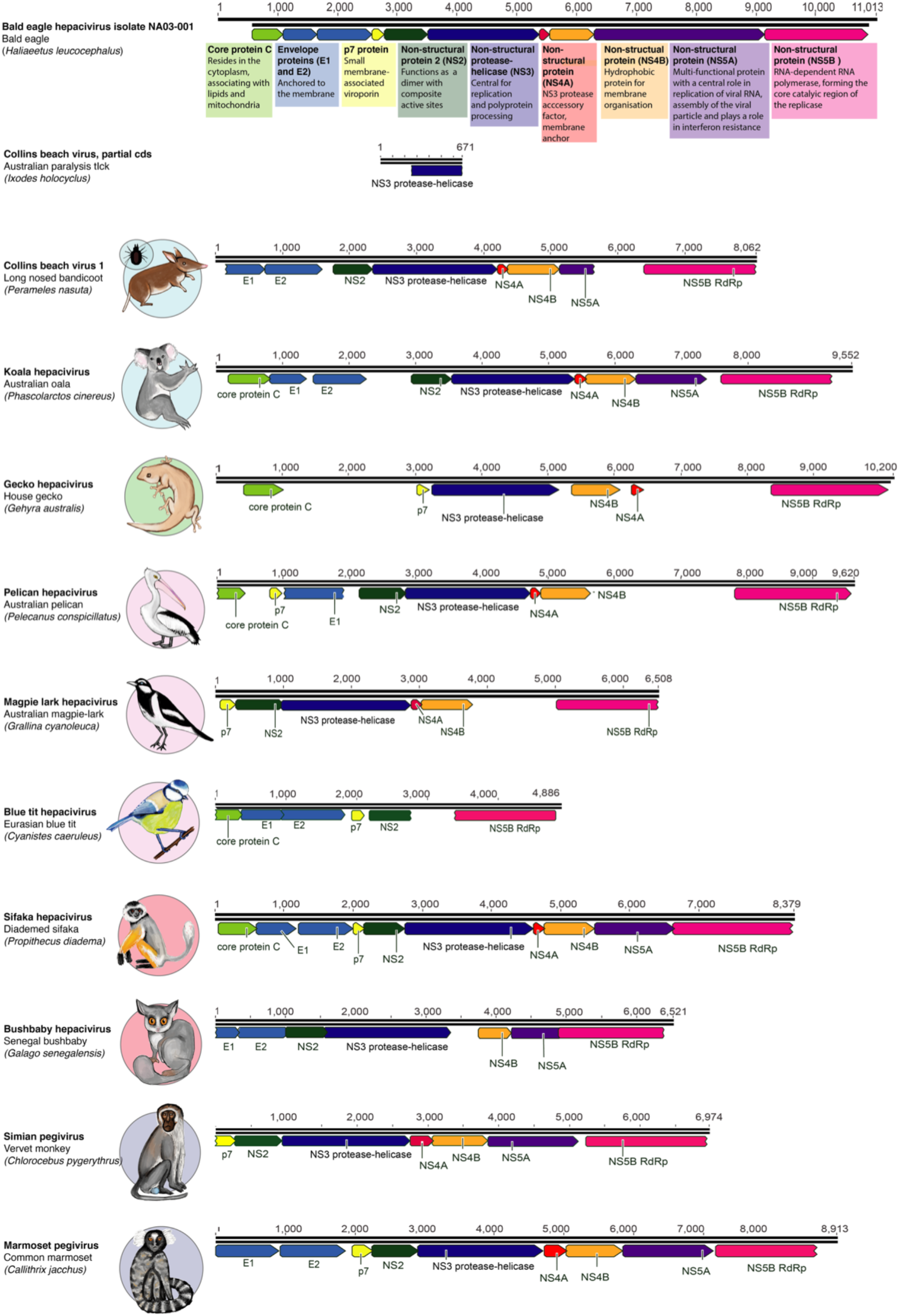
Genome annotation of the novel viruses identified in this study. The genome annotation of each virus is shown in comparison to Bald eagle hepacivirus (isolate NA03-001, accession MN062427) as a reference. The conserved *Flaviviridae* polyprotein is cleaved into several distinct protein products, represented in the diagram as coloured arrows. A short description of the function of each protein [59, 60] is shown underneath the Bald eagle hepacivirus genome, highlighted in coloured boxes that refer to each protein. Core protein C (light green), envelope proteins (light blue), p7 (yellow), NS2 (dark green), NS3 (purple), NS4A (red), NS4B (orange), NS5A (purple) and NS5B (pink).

### Phylogenetic relationships of the novel viruses

To determine the evolutionary relationships of the novel hepaci- and pegiviruses identified here we utilised the amino acid sequences of complete polypeptides from each genome in an alignment with other known pegiviruses and hepaciviruses (n = 110) (Figure 3). Maximum likelihood phylogenetic analysis revealed that the Australian Magpie lark hepacivirus (MaHV) and Pelican hepacivirus (PeHV) fall with known bird hepaciviruses, as does Blue tit hepacivirus (BtHV) (highlighted in the pink box, Figure 3). The three variants of previously identified duck hepacivirus (DuHV) [15] fall into a distinct, closely-related cluster within this clade, as do Bald eagle hepacivirus [15,16], MaHV, PeHV and BtHV, and Jogolong hepacivirus, recently isolated from a mosquito feeding on a bird host (see below) in Western Australia [20]. The existence of such an avian-specific clade is compatible with the notion that there has been some virus-host co-divergence across the hepaciviruses as a whole, particularly if the tree is rooted (as here) using the fish-associated hepacivirus that represent the most phylogenetically divergent host species [17] (Figure 3). For example, the lungfish hepacivirus falls as a sister-group to the tetrapod hepaciviruses as expected under virus-host co-divergence, and two turtle-infecting hepaciviruses, Softshell turtle hepacivirus and Chinese broad-headed pond turtle hepacivirus, are the sister-group to the mammalian hepaciviruses (highlighted in the brown box, Figure 3), although it is notable that two rodent hepaciviruses cluster anomalously with them. In this respect, it is noteworthy that the newly identified Gecko hepacivirus falls alongside other known gecko hepaciviruses - Yili teratoscincus roborowskii hepacivirus and Guangzi Chinese leopard gecko hepacivirus, both sampled in China - in a distinct gecko clade (highlighted in the green box, Figure 3).

**Figure 3.**
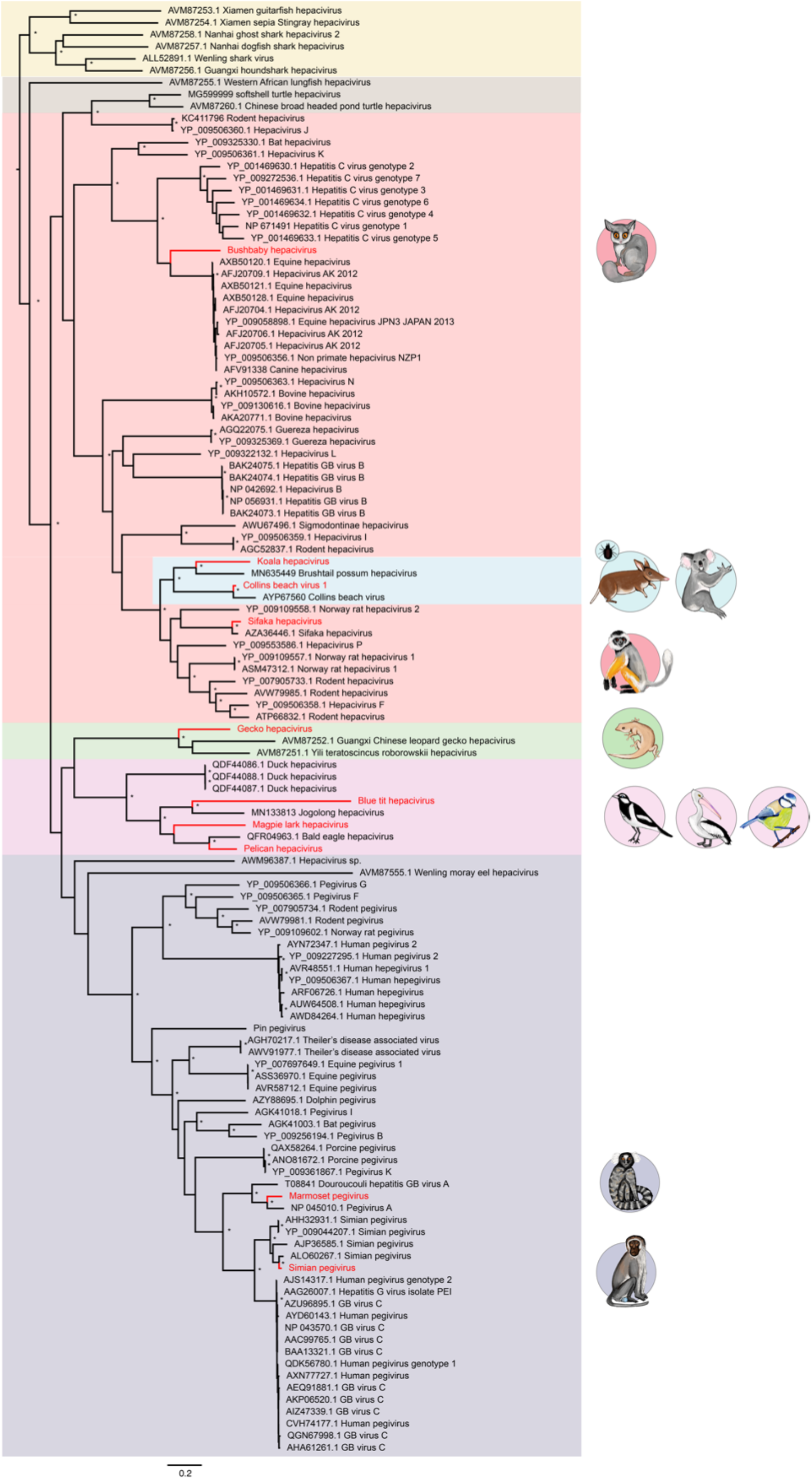
Phylogenetic analysis of the novel viruses identified in this study, identified by animal symbols, along with 106 known hepaciviruses and pegiviruses. Novel Blue tit hepacivirus, Magpie lark hepacivirus and Pelican hepacivirus, and other avian hepaciviruses are shown in the pink box. The novel Gecko hepacivirus is highlighted in the green box. Fish- and shark-associated hepaciviruses are shown in the yellow box. Turtle-associated hepaciviruses are shown in the brown box. Marsupial hepaciviruses, including those identified in bandicoots and koalas, are shown in the blue box. Bushbaby hepacivirus and Simian hepacivirus fall in the mammalian clade shown in the red box. Two novel pegiviruses, Marmoset pegivirus and Simian pegivirus, are shown in the purple box. Bootstrap values greater than 85% are shown next to the relevant nodes, represented by an asterisk. All horizontal branch lengths are scaled according to the number of amino acid substitutions per site.

The mammalian hepacivirus clade is the largest and best characterised and contains four of the novel viruses identified in this study. First, the two novel primate hepaciviruses identified via SRA mining fall within this clade, although they occupy different phylogenetic positions (Figure 3, red box). Sifaka hepacivirus (SfHV-mad) groups with a previously identified SfHV strain [61], and both viruses were isolated from different samples of *Propithecus diadema* and share 91% amino acid identity. These two viruses then cluster with a group of rodent hepaciviruses. In contrast, Bushbaby hepacivirus (BbHV), sampled from a Senegal bushbaby, groups with the equine hepaciviruses (termed NPHV, equine hepaciviruses or Hepacivirus AK) and canine hepaciviruses (CHV). Notably, sequences of the human pathogen HCV fall as a sister-group to this clade. Importantly, however, as the tree is poorly supported on the Bushbaby hepacivirus branch, it is still uncertain whether Bushbaby hepacivirus is more closely related to the equine/canine hepaciviruses or to HCV.

The remaining novel hepaciviruses identified in the mammalian clade were obtained from marsupial hosts, the Australian koala and long-nosed bandicoot, although via tick bloodmeal in the case of Collins Beach virus 1. These viruses fall in a distinct marsupial cluster that is most closely related to the hepaciviruses identified in rodents and the sifaka. In addition to these two novel marsupial hepaciviruses, the previously identified Brushtail possum hepacivirus and Collins beach hepacivirus fall into this clade. Notably, the partial sequence of Collins beach hepacivirus was found in the same pooled tick libraries as the Collins beach virus 1 described here and share relatively high sequence similarity (74% nucleotide identity, 89% amino acid identity). The ticks that made up these libraries were sampled from long-nosed bandicoots in New South Wales, Australia [19].

Two novel pegiviruses were identified from the SRA - Simian pegivirus (SPV-saf) and Marmoset pegivirus (Table 2). These fall in distinct positions within the pegivirus clade of the phylogenetic tree in a manner that seems to follow the evolutionary relationships of their hosts (Figure 3). Specifically, Simian pegivirus, identified in a Vervet monkey sample, falls close to other Old World primate-associated pegiviruses, including Simian pegiviruses isolated from Yellow baboons (*Papio cynocephalus*) from Tanzania [62], African green monkey (*Chlorocebus sabaeus*) from Gambia, and Red colobus monkey (*Piliocolobus tephrosceles*) from Uganda [63]. In contrast, Marmoset pegivirus, isolated from a common marmoset, falls with New World primate-associated pegiviruses (Pegivirus A, Douroucouli hepatitis GB virus A), identified in tamarin monkeys (*Saguinus labiatus*), mystax monkeys (*Saguinus mystax*) and owl monkeys (*Aotus trivirgatus*) [64].

Finally, it is notable two previously identified hepaciviruses, Hepacivirus sp., identified in a Red-eared slider turtle (*Trachemys scripta elegans*) and Wenling moray eel hepacivirus, identified from a moray eel (*Gymnothorax reticularis*), fall basal to all known pegiviruses, even though other fish and turtle viruses group with the hepaciviruses. To investigate this issue in more detail we performed an additional phylogenetic analysis of amino acid sequences of the NS2/3 and NS5 regions separately (Figure 4). While the phylogenetic position of the Red-eared slider turtle remains unchanged, the moray eel virus fell in markedly different positions in these two phylogenies, falling closer to the other fish hepaciviruses in the NS2/3 phylogeny (as expected with virus-host co-divergence) yet in a highly divergent position in the NS5 phylogeny. While the underlying cause of these disparate phylogenetic positions is uncertain and could plausibly have resulted from with recombination (with unknown parental sequences) or extreme rate variation, it does mean that the position of the moray eel virus as a sister-group to the pegiviruses in the polyprotein phylogeny is artifactual. While a number of other clades change position between the NS2/3 and NS5 phylogenies, particularly the fish and lungfish viruses that occupy basal positions in the polyprotein phylogeny, the deeper nodes on the NS2/3 and NS5 trees receive only weak bootstrap support such that these movements may simply reflect a lack of phylogenetic resolution. In addition, the position of the viruses newly identified here relative to those described previously remained unchanged between the NS2/3 and NS5 phylogenies.

**Figure 4.**
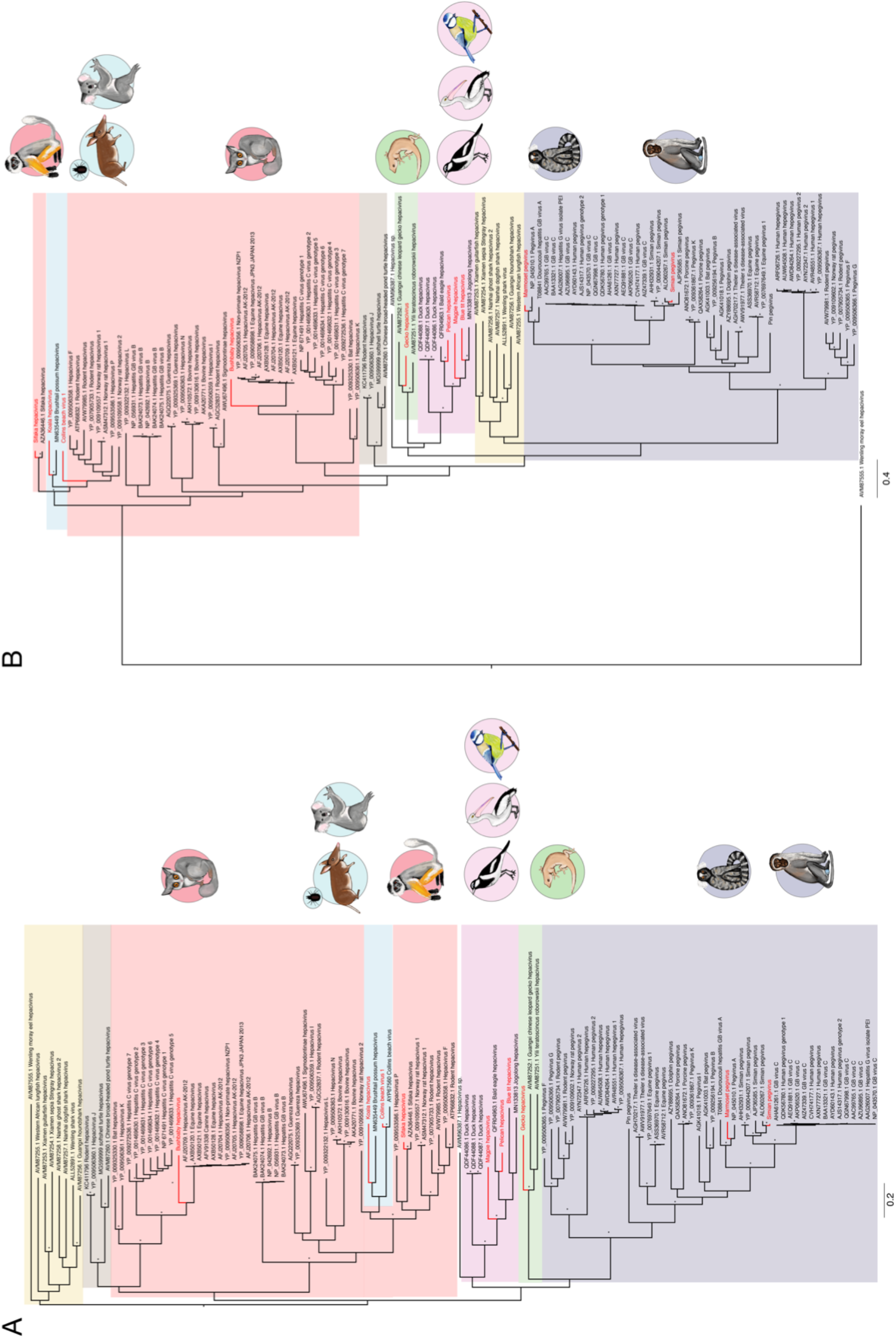
Phylogenetic analysis of the (A) NS2/NS3 and (B) NS5 amino acid sequences from the novel hepaciviruses and pegiviruses identified in this study, marked by animal symbols, combined with 106 known hepaciviruses and pegiviruses. Animal groups are shaded as described in Figure 3. Bootstrap values greater than 85% (from a total of 100 bootstrap replicates in each case) are shown next to the relevant nodes, represented by an asterisk. All horizontal branch lengths are scaled to the number of amino acid substitutions per site. Note that Collins beach virus is necessarily excluded from the NS5 phylogeny as it lacks this gene region.

## Discussion

Using a meta-transcriptomic approach we identified four novel hepaciviruses and a single variant of a previously described hepacivirus in Australian marsupials, birds, and a reptile. Additionally, through SRA-mining, we were able to discover three novel hepaciviruses and two novel pegiviruses in primate hosts. Accordingly, this work broadens our knowledge of two viral genera within the *Flaviviridae* and suggests that, with increased sampling, hepaciviruses and pegiviruses will likely be identified in many other animal hosts.

Hepaciviruses encode a single polyprotein that is commonly observed among vertebrate members of the *Flaviviridae*. The hepaciviruses identified in this study were characterised by high conservation in the NS5B, RdRp and NS3 genes. Other proteins were present in most of the viral genomes identified here, although the NS5A gene was not readily identified in the avian and gecko hepaciviruses, while the two envelope genes could not be identified in the Gecko hepacivirus. Based on the genome structure of the novel hepaciviruses, and that complete encoding region of the polyprotein is likely present, we anticipate that these genes are present in these viruses but are highly divergent in sequence and hence cannot be easily be identified using approaches that only assess primary sequence homology.

As hepaciviruses and pegiviruses are commonly thought to have co-diverged with their hosts over time-scales of many millions of years, we expected that these viruses would follow an evolutionary pattern similar to that of their hosts [17]. Indeed, our phylogenetic analysis demonstrates that this pattern generally holds true. In particular, the avian, reptile, mammal and marsupial viruses fall into distinct clades and the virus phylogeny broadly follows that of the hosts, although a number of the key nodes are only poorly supported. There are, however, some notable exceptions, such as rodent hepacivirus and Hepacivirus J, identified in bank voles (*Myodes glareolus*) from Germany [8], that clustered with turtle-associated hepaciviruses. Similarly, rather than falling as a sister-group to those hepaciviruses sampled from placental mammals as expected under virus-host co-divergence, it is notable that the marsupial hepaciviruses fall within the mammalian (eutherian) phylogenetic diversity. Finally, novel Pin pegivirus [25], isolated from a Common myna bird (*Acridotheres tristis*), falls within the diversity of mammalian pegiviruses. As this is the first avian pegivirus identified, it is possible that a distinct avian clade will be identified with increased sampling.

The avian hepacivirus clade characterised here contains Blue tit hepacivirus (BtHV), recovered from the SRA, as well the novel Australian pelican and magpie lark hepaciviruses (PeHV and MaHV, respectively), three variants of duck hepacivirus [15], and the recently described Jogalong virus, identified from a *Culex annulirostris* mosquito in northern Western Australia. However, as discussed by the authors, the mosquito from which Jogalong virus was identified is hypothesised to have recently fed upon a tawny frogmouth (*Podargus strigoides*) [20]. Similarly, a distinct clade of gecko hepaciviruses was identified, comprising the novel Gecko hepacivirus from an Australian gecko (*Gehyra lauta*)[45], and the two previously identified gecko hepaciviruses from China. The gecko clade itself clusters with the avian hepaciviruses, which in turn group with the pegiviruses. Importantly, however, there is relatively weak support for the nodes in question, and if this reptile-avian cluster in fact grouped with the hepaciviruses then the overall phylogeny would offer better support to virus-host co-divergence.

The novel Koala hepacivirus and Collins beach virus 1 (identified in ticks feeding on a long-nosed bandicoot) group closely together, along with the single marsupial-infecting hepacivirus identified previously, Brushtail possum hepacivirus [38]. Together, these viruses establish a distinct marsupial clade of hepaciviruses. Both Collins beach virus and Collins beach virus 1 also fall in this clade and we suggest that they represent variants of the same virus. Interestingly, two marsupial-associated gammaherpesviruses (double-strand DNA viruses) from Australian koalas (*Phascolarctos cinereus*) and wombats (*Vombatus ursinus*), similarly formed a distinct clade [65]. Hence, marsupial-specific clades may be present in other vertebrate-infecting viruses and undergo co-divergence with their hosts. Additionally, two primate hepaciviruses were identified from the SRA, including a novel variant of SfHV (Sifaka hepacivirus) closely related to the previously identified SfHV, both of which were isolated from Diademed sifakas. In contrast, Bushbaby hepacivirus isolated from a Senegal bushbaby, instead falls as a sister-group to a cluster of equine and canine hepaciviruses that are in turn closely related to HCV. Despite the obvious clinical importance of HCV, its ultimate animal reservoir is unknown [22–25]. The placement of another primate (i.e. bushbaby) virus in this clade is notable as it suggests that there are additional primate viruses within this group that are yet to be discovered and that might shed light on the origin of HCV.

Primate pegiviruses fall into distinct clades – in Old World versus New World primates - and the two novel sequences identified here follow this pattern. Several pegiviruses have been detected in a variety of Old World monkeys, including red colobus monkeys (*Procolobus ephroscles*), red-tailed guenons *(Cercopithecus ascanius)* and an olive baboon (*Papio Anubis)* [62, 63], as well as the novel Simian pegivirus genome identified here in a South African vervet monkey. Similarly, the New World pegiviruses form a distinct clade that includes Pegivirus A (also termed GB virus A), isolated from tamarins [64, 66, 67], and newly identified Marmoset pegivirus, isolated from a Common marmoset.

Arthropod vectors are known to transmit a wide variety of infectious disease agents, and are responsible for more than 17% of total infectious diseases and cause upwards of 700,000 deaths per year [68]. Notably, the *Flaviviridae* contain many arthropod-borne viruses that impose a serious burden on human populations, including yellow fever virus, Zika virus, and dengue virus. However, all the vector-borne flaviviruses described to date are members of the genus *Flavivirus*, with no evidence of vector-borne transmission within the hepaciviruses or pegiviruses. Although the data presented here and previously [19] tentatively suggest that *I. holocyclus* ticks may act as a vector of Collins beach virus, the biology of ticks prevents us from clearly establishing this link [69]. In particular, as the ticks sampled for RNA extraction were engorged adult females it is impossible to determine if Collins beach virus infected the tick itself or was merely held within their considerable blood meal. As ticks take only one blood meal in each lifecycle stage (of which adulthood is the final stage) they do not act as ‘biological syringes’ as other arthropods such as mosquitos and biting flies do [70]. As Collins beach virus and Collins beach virus 1 were detected in very low abundance it does not seem likely that this virus is actively infecting the tick itself. Future meta-transcriptomic studies on unfed questing ticks are therefore a priority.

In sum, we have expanded the diversity of known hepaciviruses and hypothesise that a wide range of hosts that are yet to be identified globally. Unsurprisingly Australia’s unique fauna host an equally diverse virome. The meta-transcriptomic tools described here can help us to explore the breadth and depth of viral assemblages, investigate the role of putative pathogens in wildlife disease syndromes, and contribute to the rapid and accurate diagnosis of emergent disease syndromes.

## Supplementary Materials

**Supplementary Figure 1**. Individual fragments used in compiling the Blue tit hepacivirus and Simian pegivirus genomes. Each fragment has a coding region with draft annotations of key viral proteins. (A) The three fragments from SRA project SRX565268 (SRR1325073, SRR1325072, SRR1325074) from the Eurasian blue tit *(Cyanistes caeruleus)* from Germany. (B) The two fragments (SRR1046735, SRR1046733) from SRA project SRX389659, taken from a Vervet monkey (*Chlorocebus pygerythrus*) sampled in South Africa. These two fragments were concatenated into a single genome, Simian pegivirus (SPV-saf).

**Supplementary Table 1.** Results of the SRA mining of marsupial transcriptomes.

**Supplementary Table 2.** Abundance of hepacivirus reads in the total read count of the sequencing libraries used in this analysis.

**Supplementary Table 3.** Results of the SRA mining of avian transcriptomes (excluding *Gallus gallus*).

**Supplementary Table 4.** Results of the SRA mining of primate transcriptomes (excluding *Homo sapiens*).

**Supplementary Table 5.** GenBank accession numbers and description of the hepacivirus and pegivirus amino acid sequences used in the phylogenetic analysis.

## Author Contributions

Conceptualization, E.C.H; Methodology, E.C.H, J.H.O.P., J-S.E, J.P.B, C.M., K.R, A.F.P; Formal Analysis, A.F.P, J.H.O.P., J-S.E; Writing – Original Draft Preparation, A.F.P, E.C.H; Writing – Review & Editing, A.F.P, E.C.H, E.H.,J.H.O.P., W-S.C, J-S.E; Funding Acquisition, E.C.H.

## Data Availability

The sequences have been submitted to GenBank with accession numbers MT371434-MT371443.

## Acknowledgments

Analysis was performed using the University of Sydney Artemis HPC. We thank the Australian Museum for their generous provision of sample material used in this study.

