## Supplementary figures and images for "Metagenomic identification of diverse animal hepaciviruses and pegiviruses"

### Supplementary Figure 1

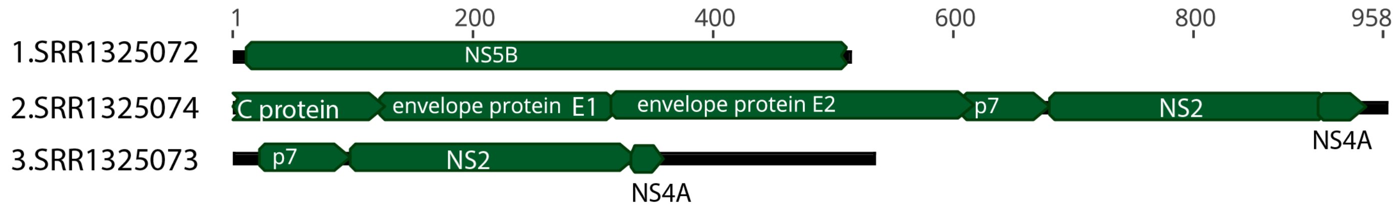

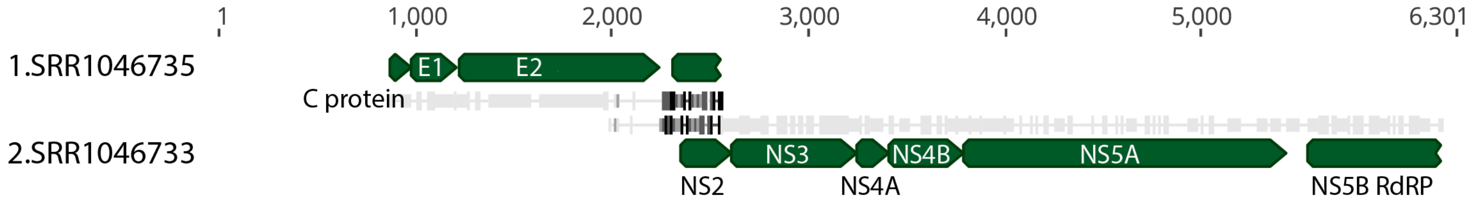
